# Somatodendritic release of cholecystokinin potentiates GABAergic synapses onto ventral tegmental area dopamine cells

**DOI:** 10.1101/2022.01.14.476405

**Authors:** Valentina Martinez Damonte, Matthew B. Pomrenze, Caroline Casper, Annie M. Wolfden, Robert C. Malenka, Julie A. Kauer

## Abstract

Neuropeptides are contained in nearly every neuron in the central nervous system and can be released from somatodendritic sites as well as from nerve terminals. Cholecystokinin (CCK), among the most abundant neuropeptides in the brain, is expressed in the majority of midbrain dopamine neurons. Here we report that ventral tegmental area (VTA) dopamine neurons release CCK from somatodendritic regions, where it triggers long-term potentiation of GABAergic synapses. The somatodendritic release occurs with trains of action potentials or prolonged but modest depolarization and is dependent on synaptotagmin 7 and T-type Ca^2+^ channels. Depolarization-induced LTP is blocked by the CCK2R antagonist, LY225910, and mimicked by exogenously added CCK. To test the behavioral role of CCK, we infused it into the mouse VTA. Ca^2+^ imaging *in vivo* demonstrated that infused CCK reduced dopamine cell signals during fasted food consumption. Moreover, local infusion of CCK also inhibited food consumption and decreased distance traveled in an open field test. Together our experiments introduce somatodendritic neuropeptide release as a previously unknown feedback regulator of VTA dopamine cell excitability and dopamine-related behaviors.

## INTRODUCTION

Nearly every neuron in the brain contains neuropeptides (Smith et al., 2020). Acting at G-protein coupled receptors, they can regulate neuronal excitability by modulating voltage-gated ion channels and can modify synaptic transmission mediated by classical neurotransmitters (Ludwig and Leng, 2006; Smith *et al*., 2020). Neuropeptides are synthesized in somata and dendrites and are then cleaved and stored in large dense core vesicles (LDCV), which are axonally transported and released by exocytosis from nerve terminals. However, neuropeptides are also released at somatodendritic sites (Brown et al., 2020; Crosby et al., 2015; Crosby et al., 2018; Krawczyk et al., 2013; Ludwig and Leng, 2006; Wagner et al., 1993). In fact, dense core vesicles that contain peptides are found in similar numbers in axons and dendrites, (Lipka et al., 2016; Persoon et al., 2018; Zahn et al., 2004). The best-studied examples of somatodendritic peptide release are oxytocin and vasopressin, which serve autocrine and paracrine roles in regulating hypothalamic neuron excitability (Brown *et al*., 2020; Ludwig and Pittman, 2003). Somatodendritic peptide release may thus serve functional roles as important as neuropeptide release from nerve terminals.

Originally identified in the gut as a modulator of digestive function, cholecystokinin (CCK) is one of the most highly expressed neuropeptides in the brain (Crawley, 1985; Rehfeld, 2017). CCK is co-expressed with dopamine (DA) in the majority of ventral tegmental area (VTA) neurons in the rat, monkey and mouse (Crawley and Corwin, 1994; Heymann et al., 2020; Hokfelt et al., 1986; Hokfelt et al., 1980; Jayaraman et al., 1990; Seroogy et al., 1988). In rats, exogenous CCK can transiently excite VTA dopamine cells (Brodie and Dunwiddie, 1987; Skirboll et al., 1981; Stittsworth and Mueller, 1990) but it also potentiates dopamine-induced inhibition of DA cell firing in rat VTA *in vitro* and *in vivo* (Artaud et al., 1989; Brodie and Dunwiddie, 1987; Chiodo et al., 1987). Although the first example of somatodendritic release was dopamine from DA cells (Cheramy et al., 1981), somatodendritic neuropeptide release from these neurons has not been described.

Controlling the excitability of midbrain dopamine neurons is critical since changes in DA release shape multiple physiological phenomena including motivation, motor function and learning (Berke, 2018; Berridge, 2007; Cox and Witten, 2019; Friedman, 2014; Howe and Dombeck, 2016; Popescu et al., 2016; Salamone and Correa, 2012). Both local and extrinsic GABAergic afferents control the firing of DA neurons, and thus plasticity of inhibitory synapses has a major influence on these circuits (Beckstead and Williams, 2007; Polter and Kauer, 2014; Simmons et al., 2017; Xin et al., 2016). We previously reported that inhibitory GABAergic synapses in the VTA undergo long-term potentiation (LTP) following afferent stimulation paired with depolarization (St Laurent and Kauer, 2019; St Laurent et al., 2020). These studies indicated an NMDAR-independent mechanism for the LTP that requires depolarization of the postsynaptic dopamine cell and is blocked by postsynaptic chelation of Ca^2+^, features consistent with somatodendritic release of a neuromodulator that mediates LTP induction. Here we report that somatodendritic release of CCK underlies LTP of specific GABAergic synapses onto VTA dopamine neurons. We also demonstrate that CCK depresses VTA DA cell activity, feeding, and locomotor behaviors. Together our work suggests that CCK released somatodendritically from DA neurons will exert potent behavioral effects.

## RESULTS

### Depolarization of a postsynaptic dopamine neuron potentiates GABA_A_ IPSCs

We previously reported that low frequency stimulation (LFS) paired with sustained modest depolarization induces LTP at GABA_A_ synapses in the VTA (St Laurent and Kauer, 2019)(Figure 1A-C), but the mechanisms were unclear. We hypothesized that even in the absence of LFS, the depolarization could release a signaling molecule from the recorded DA neuron that in turn elicits LTP. To test whether depolarization alone could potentiate GABAergic IPSCs, we recorded IPSCs evoked with a stimulating electrode placed caudal to the VTA in horizontal slices. GABAergic IPSCs were measured before and after 6 min of depolarization to -40 mV, without concomitant synaptic stimulation. This maneuver caused LTP of a magnitude similar to that triggered by LFS plus depolarization (Figure 1D-F). Thus, LFS is not required, but depolarization alone is sufficient to initiate LTP. To further test whether depolarization of DA cells is sufficient to induce LTP, we took an orthogonal approach by expressing channelrhodopsin (ChR2) in DAT-Cre mice and stimulating DA cells with blue light. Optogenetically driving DA cells to fire repetitively also triggered LTP of evoked GABAergic IPSCs (Figure 1G-I).

**Figure 1.**
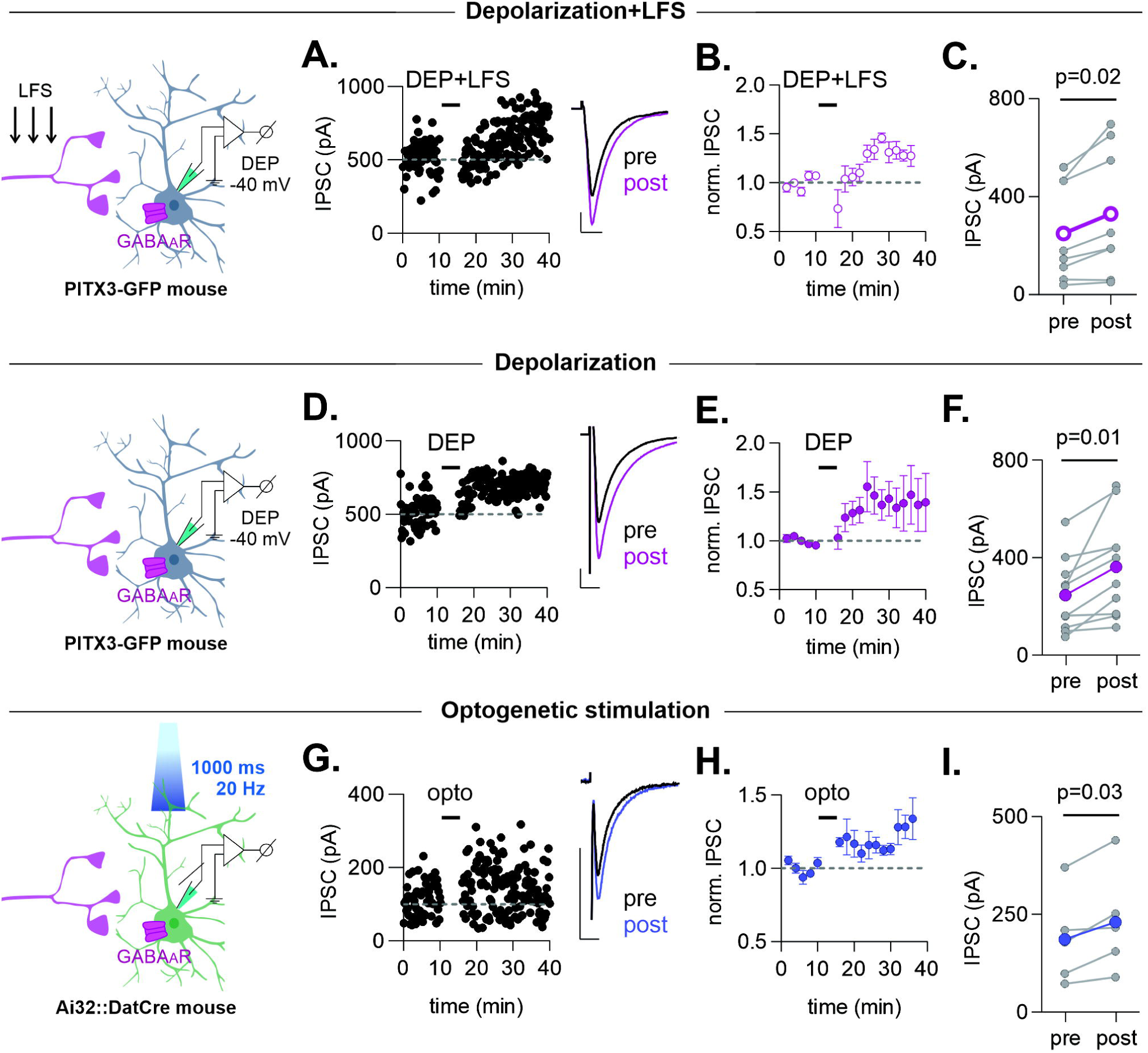
Depolarization alone potentiates inhibitory postsynaptic currents (IPSCs) in VTA dopamine cells. Left-hand diagrams in this and all figures illustrate the experimental design. **A.** Representative time course and example IPSCs (inset) before and after a 6-minute depolarization of the recorded dopamine neuron from -70 to -40 mV with simultaneous afferent low frequency (1 Hz) stimulation (DEP+LFS). **B.** Averaged IPSC amplitudes before and after DEP+LFS (n=8 cells/8 mice). **C.** IPSC amplitudes before and after DEP+LFS (n=8 cells/8 mice). Wilcoxon t-test, p=0.02. For this and all figures, colored symbols/lines represent the mean. Error bars represent SEM. **D.** Representative time course and example IPSCs before and after a 6-minute depolarization of the recorded neuron from -70 to -40 mV (DEP). Data for 1A-D are from a wild type mouse. **E.** Averaged IPSC amplitudes before and after DEP (n=10 cells/8 mice). **F.** IPSC amplitudes before and after DEP (n=10 cells/8 mice). Paired t-test, p=0.01. **G.** Representative time course and example IPSCs from a VTA dopamine neuron expressing ChR2 (Ai32::Dat-Cre mouse,) before and after optical stimulation; trains of light (20 Hz, 1000 ms) were delivered for 6 minutes in current clamp. **H.** Time course of averaged IPSC amplitudes before and after optical stimulation (n=5 cells/ 3 mice). **I.** IPSC amplitudes before and after optical stimulation (n=5 cells/3 mice). Paired t-test, p=0.03. Data for 1E-I are from Pitx3-GFP labeled dopamine neurons. Scale bars, 100 pA,10 ms.

Dopamine has long been known to be released somatodendritically from midbrain DA neurons and is therefore a candidate to mediate LTP at GABAergic nerve terminals via activation of one or both dopamine receptor classes. To investigate this possibility, we recorded IPSCs while blocking both D1 and D2 dopamine receptors. Application of sulpiride (150 nM) and SCH 23390 (10 µM) did not prevent depolarization-induced LTP, indicating that DA release is not involved in this form of LTP (Figure S2A-C).

### DA neurons contain CCK and release it with optogenetic stimulation

GFP-positive cells from Pitx3-GFP mice have ∼95% concordance with DA neurons in the VTA (Maxwell et al., 2005)). Using these mice, we confirmed that CCK is highly expressed in VTA DA cells (Figure S1A-C) (75.3 ± 1.8% GFP-positive neurons were also CCK-positive). Consistent with this finding, we found a 77.5 ± 1.6% overlap of mRNA for DAT and CCK using FISH (Figure S1 D-F).

It was previously shown that in rat midbrain slices high K^+^ stimulation evoked CCK release measured by radioimmunoassay, but this protocol depolarizes the entire preparation and is expected to release CCK from both somatodendritic and axon terminal sites (Freeman et al., 1991). In a more targeted approach, we used slices from DAT-Cre mice selectively expressing ChR2 in DA cells, stimulated with blue light, and measured CCK with an ELISA assay. The CCK concentration in the supernatant increased after optogenetic stimulation (1.3 ±0.9 pg/ml vs 6.3 ±1.6 pg/ml) demonstrating that CCK can be released by optogenetic excitation of DA neurons.

### LTP is blocked by a CCK2R antagonist

Might CCK release occurring during depolarization cause LTP? CCK acts via two G-coupled receptors, CCK1R and CCK2R (Dufresne et al., 2006; Hill and Woodruff, 1990), and the CCK2R is predominant in the brain. We therefore bath-applied the CCK2 receptor antagonist, LY225910, and tested whether depolarization was still capable of inducing LTP. Neither depolarization alone (Figure 2A-C) nor depolarization paired with LFS (Figure S2D-F) potentiated IPSCs in the presence of the CCK2R antagonist. Furthermore, using FISH, we did not detect expression of CCK1R (data not shown) or CCK2R mRNA within the VTA (Figure 2D), indicating that these receptors are not found in somata or glial cells within the VTA. These results suggest that CCK acting via the CCK2R is necessary for LTP, and that CCK2Rs are likely to be localized on afferent nerve terminals rather than in the VTA cell population.

**Figure 2.**
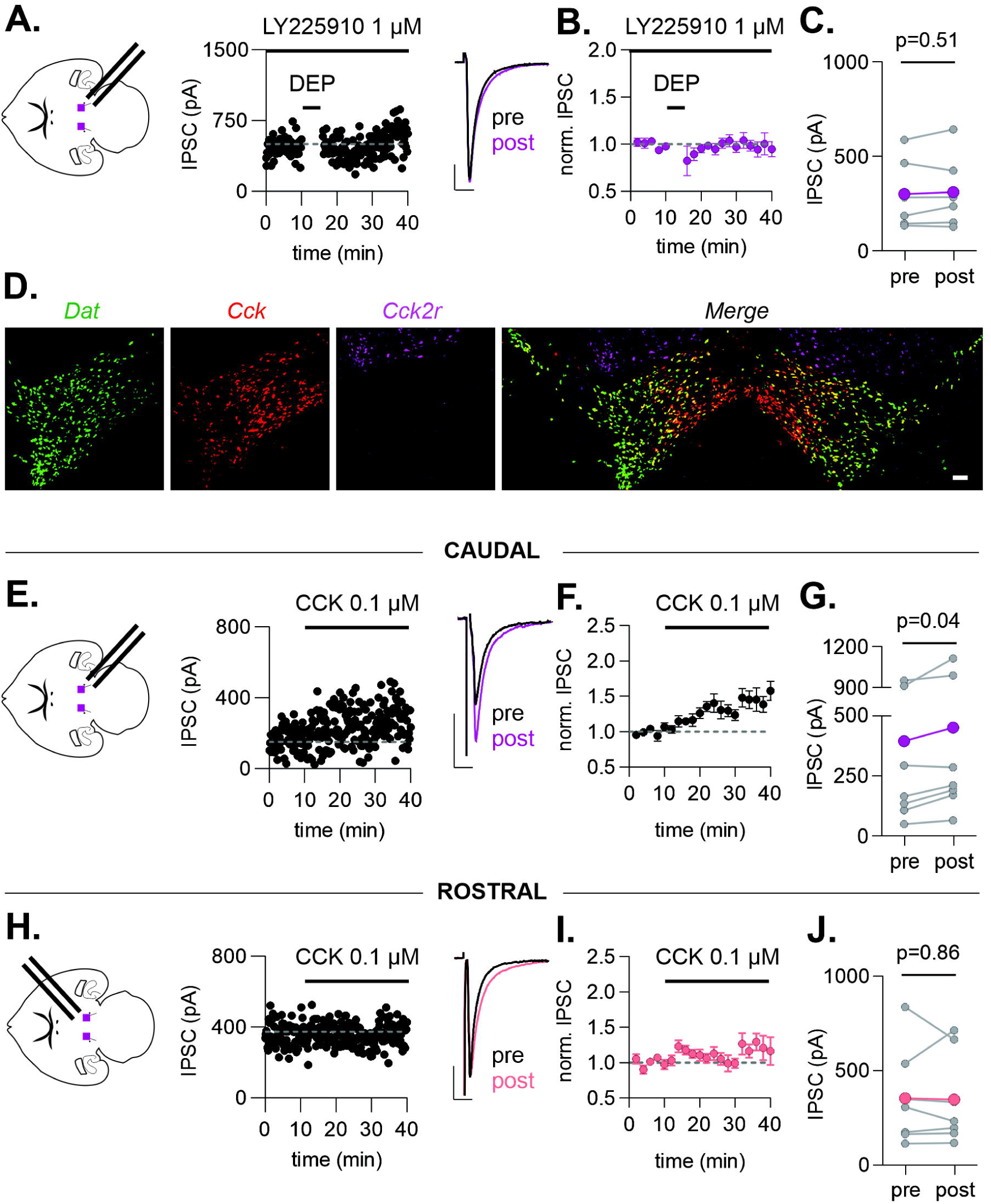
CCK is necessary and sufficient to potentiate caudally-evoked but not rostrally-evoked IPSCs. **A.** Representative time course and example IPSCs before and after a 6-minute depolarization of the recorded dopamine neuron from -70 to -40 mV (DEP) in the presence of the CCK2R antagonist, LY225910 (1 µM). **B.** Averaged IPSC amplitudes before and after DEP (n=6 cells/3 mice). **C.** IPSC amplitudes before and after DEP (n=6 cells/3 mice). Paired t-test, p=0.51. **D.** Representative confocal images of a FISH experiment on a coronal slice containing the VTA. *Dat* (green), *Cck* (red), *Cck2r* (magenta) and Merge: overlay of all three signals. **E.** Representative time course and example IPSCs evoked in a dopamine neuron with a stimulating electrode placed caudal to the VTA (diagram) before and during bath application of CCK (0.1 µM). **F.** Averaged caudally-evoked IPSC amplitudes before and during CCK (n=8 cells/7 mice). **G.** Caudally-evoked IPSC amplitudes before and after CCK (n=8 cells/7 mice). Paired t-test, p=0.04. **H.** Representative time course and example IPSCs evoked in a dopamine neuron with a stimulating electrode placed rostrally within the VTA (diagram) before and after bath application of CCK (0.1 µM). **I.** Averaged rostrally-evoked IPSC amplitudes before and after CCK (n=7 cells/5 mice). **J.** Rostrally-evoked IPSC amplitudes before and after CCK (n=7 cells/5 mice). Paired t-test, p=0.86. Scale bars, 100 pA,10 ms.

### CCK occludes LTP of IPSCs and acts in a synapse-specific manner

If depolarization-induced CCK release from a single recorded DA neuron is sufficient to trigger LTP, then application of exogenous CCK should mimic this. Consistent with this prediction, bath application of CCK elicited LTP with a time course and of a magnitude similar to LTP evoked with depolarization (Figure 2E-G). Moreover, after CCK application, depolarization did not elicit further LTP, arguing for a shared mechanism (Figure S2G-I). To this point, all our experiments used a stimulating electrode positioned caudal to the VTA. Because previous studies reported that depolarization does not potentiate rostrally-evoked IPSCs (Dacher and Nugent, 2011; St Laurent and Kauer, 2019), we next asked whether CCK-induced potentiation also exhibits synapse-specificity. Consistent with these previous results and in contrast to the observed CCK-induced LTP of caudally-evoked IPSCs, IPSCs stimulated with a rostrally-placed stimulating electrode were unaffected by CCK (Figure 2H-J). Together our data demonstrate that CCK is necessary and sufficient for selective potentiation of caudally-evoked IPSCs.

### CCK does not modulate D2-IPSCs

The coexpression of CCK and DA and their shared ability to undergo somatodendritic release from DA neurons prompted us to investigate their interplay. Train-evoked postsynaptic inhibitory currents mediated by somatodendritic release of dopamine acting at D2 autoreceptors (Beckstead et al., 2004) are a convenient readout of somatodendritic DA release. To test the possibility that CCK modifies somatodendritic DA release, we simultaneously evoked D2-IPSCs and caudal GABAergic IPSCs in the same cells, and bath applied CCK. While CCK potentiated GABAergic IPSCs as expected, it had no effect on D2-IPSCs (Figure 3). Thus, CCK does not appear to influence somatodendritic release of dopamine as assayed by measurement of the D2-IPSC.

**Figure 3.**
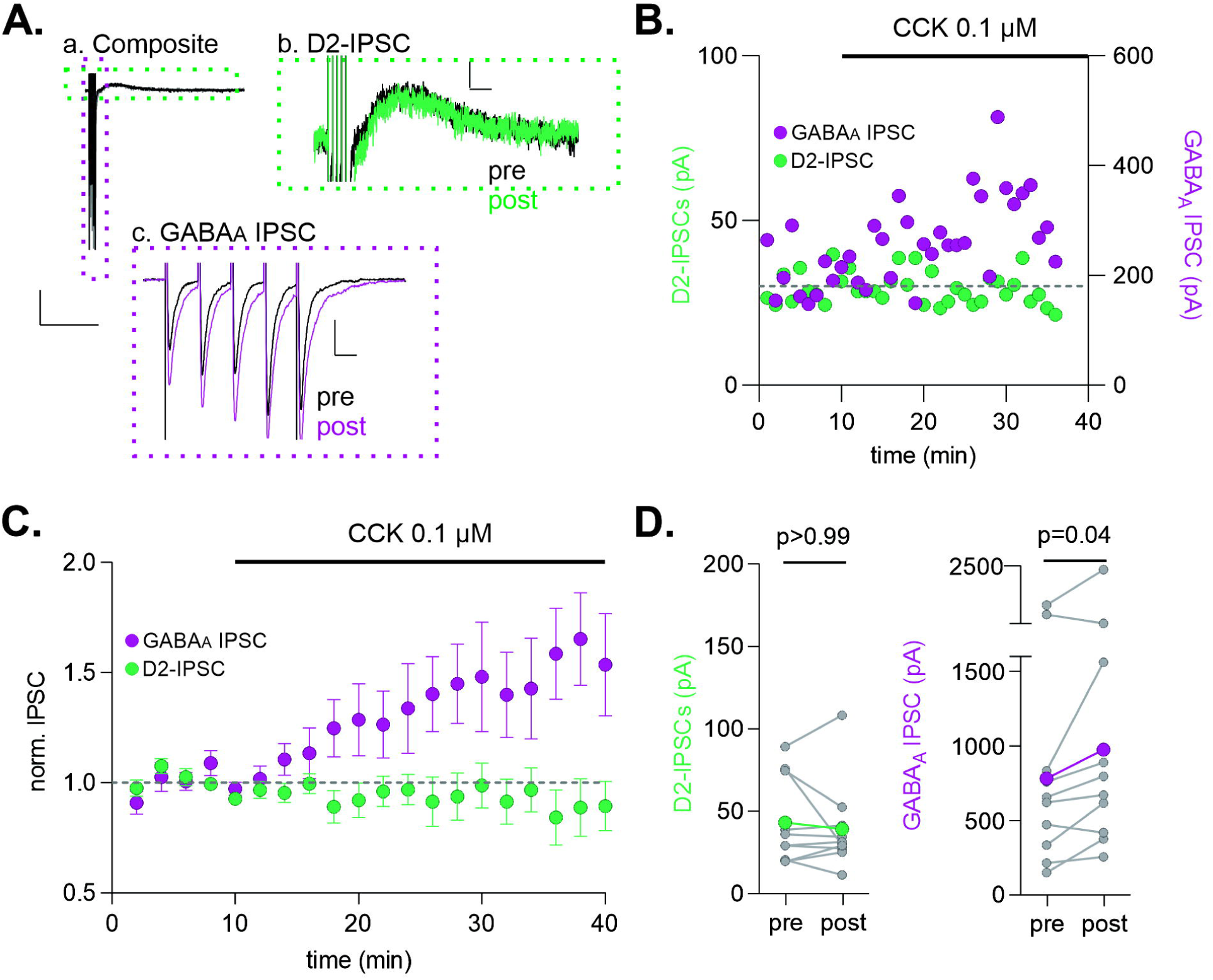
CCK potentiates GABA_A_ synapses but does not affect D2-IPSCs. **A.** Representative examples of (a) composite IPSCs, (b) D2-IPSCs, and (c) GABA_A_ IPSCs recorded in a single dopamine neuron before and after CCK (0.1µM) bath application. **B.** Representative example time course of simultaneously recorded D2-IPSCs (green) and GABA_A_ IPSCs (purple). **C.** Averaged IPSC amplitudes before and after CCK bath application (n=10 cells/9 mice). **D.** D2-IPSC and GABA_A_ IPSCs amplitudes before and after CCK (n=10 cells/9 mice). Wilcoxon t-test: D2-IPSCs, p>0.99); GABA_A_ IPSCs, p=0.04. Scale bars, (a) 100 pA,1s (b) 10 pA,10 ms (c) 100 pA,10 ms.

### CCK release requires T-type calcium channels and synaptotagmin 7

A rise in intracellular calcium is generally required for somatodendritic release of peptides, but the calcium sources involved can be diverse (Ludwig et al., 2016). Consistent with a critical role for intracellular Ca^2+^, inclusion of the calcium chelator BAPTA in the recording pipette prevented depolarization-induced LTP (Figure S3A-C). All voltage-gated calcium channel (VGCC) subtypes are expressed in DA neurons, however T-type VGCCs display ‘window currents’ in which tonic influx of calcium can occur at membrane potentials close to rest (Cain and Snutch, 2010; Evans et al., 2017; Perez-Reyes, 2003; Tracy et al., 2018). Therefore, we next tested NiCl_2_ at a concentration that primarily blocks T-type calcium channels (50 µM) (Lee et al., 1999; Perez-Reyes, 2003; Zamponi et al., 1996). Depolarization did not potentiate IPSCs in the presence of NiCl_2_, suggesting a crucial requirement for T-type channels. In contrast, D2-IPSCs could still be evoked (n= 4 of 4 cells) (Figure S3D-F).

The release machinery required for somatodendritic release of neurotransmitters differs from that at nerve terminal active zones (Ludwig *et al*., 2016). Synaptotagmins (syts) 4 and 7 have been implicated in somatodendritic release of neuropeptides as well as dopamine (Delignat-Lavaud, 2021; Mendez et al., 2011; Zhang et al., 2009). To test a role for syt7, we used syt7 knock out mice (syt7 KO mice) and found that depolarization-induced LTP was strongly attenuated in syt7 KO slices (Figure S3G-I). Moreover, in agreement with a previous report using knockdown syt 7 (Mendez *et al*., 2011), D2 IPSCs in syt7 KO mice were also markedly reduced (6/7 DA cells tested). Our results suggest that both somatodendritic DA and CCK release require syt7.

### Intra VTA delivery of CCK reduces food intake and locomotion, and DA neuron activity

What physiological processes might be controlled by the local release of CCK within the VTA? CCK-expressing VTA neurons project most strongly to the nucleus accumbens shell (Heymann *et al*., 2020; Poulin et al., 2018) and contribute to the regulation of reward behaviors, including Pavlovian and operant conditioning for sucrose rewards (Heymann *et al*., 2020). We therefore asked whether CCK infused into the VTA modulates feeding behavior. Wild-type mice were prepared with bilateral guide cannulas targeting the VTA and after overnight food deprivation, either CCK or saline was delivered locally into the VTA. Consumption of standard chow was measured over a 1-hr period. At least 5 days later the opposite compound (saline or CCK) was infused into the VTA after a second fast in a crossover design. Intra-VTA delivery of CCK reduced food intake when compared with intra-VTA delivery of saline in the same animals (Figure 4A-B; S4B). Total distance traveled over 30 minutes was also reduced in mice treated with CCK compared with saline (Figure 4C), however the modest reduction in locomotion is unlikely to account for the reduction in food intake over a 1 h feeding session.

**Figure 4.**
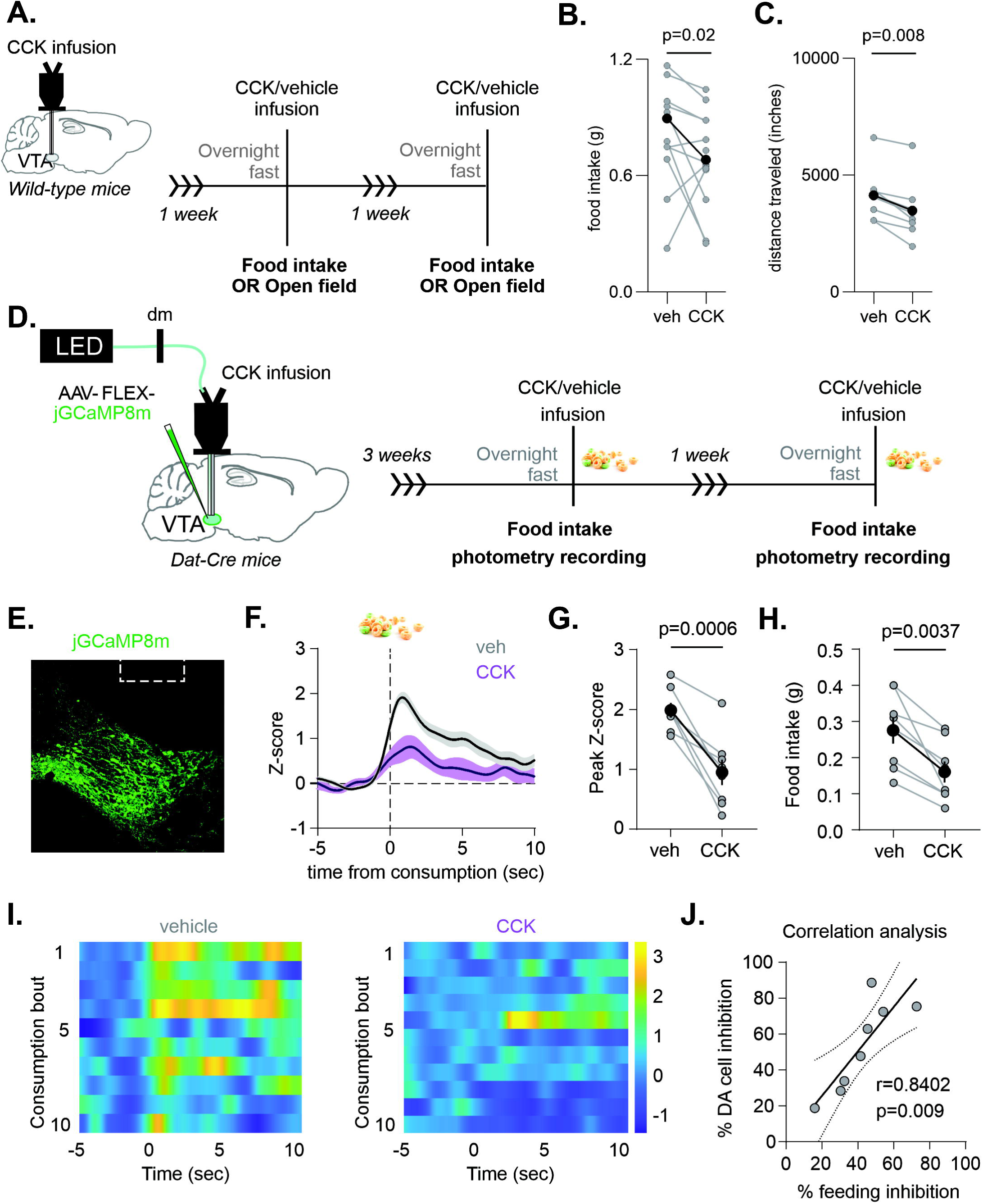
Intra-VTA CCK infusion reduces food intake and DA cell activity. **A.** Schematic diagram of experimental design for food intake or open field locomotion. **B.** Food intake over the 1-hour period after bilateral intra-VTA infusion of either CCK or vehicle (n=13 mice). Paired t-test, p=0.02. **C.** Total distance traveled in the open field apparatus over a 30 min period after bilateral intra-VTA infusion of CCK or vehicle (n=8 mice). Paired t-test, p=0.008. **D.** Schematic of fiber photometry configuration and experimental timeline. **E.** Representative image of GCaMP8m in VTA DA neurons, coronal slice. **F.** Peri-stimulus histogram of time course of averaged GCaMP8m transient z-scores event-locked to food consumption (n=8). **G.** Quantification of peak z-scores during food consumption after either saline or CCK infusion (n=8). Paired t-test, p=0.0006. **H.** Quantification of food intake during recording sessions over a 30 min period (n=8). Paired t-test, p=0.004. **I.** Representative heat map of z-score changes over all trials from individual mice; zero time is the onset of food consumption. **J.** CCK-induced DA cell inhibition and food intake are significantly correlated (n=8). Pearson’s r=0.84. Paired t-test, p=0.009. Bold symbols/lines represent the mean response across all animals, error bars represent SEM.

CCK may inhibit feeding behavior by decreasing DA cell excitability. To test this, we measured Ca^2+^ transients while mice engaged in home cage feeding. DAT-Cre mice were injected with AAV-FLEX-GCaMP8m, implanted with a cannula/optic fiber into the VTA and trained to consume palatable food (fruit loops). After an overnight fast, either CCK or saline was microinjected into the VTA, and imaging began. Following a 5 min baseline, fruit loops were presented and mice were recorded for 30 min. At least 5 days later, the alternate compound was delivered into the VTA for a second recording/feeding session in a crossover design (Figure 4D). With a saline infusion, DA cells displayed transient increases in activity time-locked to food consumption. However, after CCK DA cell activity was significantly dampened upon food consumption (Figure 4F-G and I). The amount of food consumed was also significantly reduced when CCK was infused (Figure 4H), and the degree of CCK-induced DA cell inhibition and food intake reduction (compared to magnitudes measured on saline) were highly correlated (Figure 4J). Together, these data suggest that CCK signaling in the VTA reduces DA cell activity to suppress food intake, and indicate a likely role for somatodendritically-released CCK.

## DISCUSSION

The CNS signaling roles of neuropeptides are not well understood despite their broad distribution (Smith *et al*., 2020). The expression of CCK in VTA DA neurons is a good example, as it was identified over forty years ago (Hokfelt *et al*., 1980), but even the deployment of modern tools has focused on CCK as a useful marker of cell groups rather than elucidating the functions of CCK itself (Heymann *et al*., 2020; Poulin *et al*., 2018). Here we report that CCK is released somatodendritically when DA cells are depolarized, and in turn potentiates inhibitory synapses. Our results support the idea that locally-released CCK is part of an inhibitory feedback mechanism, acting synergistically with somatodendritic DA release to inhibit DA neuron excitability and modulate dopamine-dependent behaviors. Given the widespread expression of CCK in other brain areas, this may be a general mechanism to regulate synaptic strength.

### CCK is somatodendritically released from DA cells

The CCK release we observe is likely predominantly of somatodendritic origin rather than from nerve terminals. First, only a small subset of dopaminergic nerve terminals in the VTA originate from VTA/SNc (Bayer and Pickel, 1990; Deutch et al., 1988). Second, we evoked CCK release with subthreshold depolarization of a single dopamine cell, a stimulus unlikely to trigger nerve terminal release. Finally, blocking T-type VGCCs with Ni^2+^ blocks the CCK LTP, but is not expected to affect nerve terminal release of neurotransmitter (Dolphin and Lee, 2020).

Optogenetic depolarization of DA neurons in the VTA triggers somatodendritic CCK release, which has not been accounted for in optogenetic studies to date. Moreover, other neuropeptides are expressed in DA neurons, and depolarization may trigger their release as well (Dore et al., 2020; Perez-Bonilla et al., 2021). Somatodendritic peptide release will need to be considered in studies using optogenetic activation of cell body and dendritic sites.

### Somatodendritic CCK release requires T-type Ca^2+^ channels and syt7

We find that CCK release is blocked by chelation of calcium in the DA cell, consistent with our previous work (St Laurent *et al*., 2020) and with the reported requirement for extracellular Ca^2+^ in high K^+^-induced CCK release from midbrain slices (Freeman *et al*., 1991). Among VGCCs, T-type channels are likely candidates to contribute to depolarization-induced CCK release based on their voltage-dependence (Evans *et al*., 2017; Tracy *et al*., 2018), and our data with NiCl_2_ support this hypothesis (Wolfart and Roeper, 2002). It is somewhat surprising that a 6-minute depolarization to -40mV releases CCK since DA neurons rest at relatively depolarized membrane potentials, often firing spontaneously *in vivo* and *in vitro*. However, trains of action potentials also release CCK, as optogenetic stimulation of VTA DA neurons expressing ChR2 elicited LTP (Figure 1G-H) and allowed biochemical detection of CCK release.

The Ca^2+^ sensing protein syt1 is likely to account entirely for fast DA release from striatal nerve terminals (Banerjee et al., 2020; Lein et al., 2007; Mendez *et al*., 2011; Saunders et al., 2018). However, even in DA axon fields in the striatum, high K^+^- and action potential-dependent DA release still occurs in the syt1 KO mouse, supporting the idea that another Ca^2+^ sensor, hypothesized to be syt7, is responsible for an additional slower form of release (Banerjee *et al*., 2020). Knockdown of syt7 was also reported to decrease somatodendritic DA release from cultured neurons (Delignat-Lavaud et al., 2021; Kissiwaa et al., 2021; Mendez *et al*., 2011). In our hands, both CCK release (using depolarization-induced LTP as a proxy for CCK release) and the D2-IPSC (a readout of somatodendritic DA release) were absent in the syt7 KO mouse. Our data support a model in which Ca^2+^ entry through T-type Ca^2+^ channels, perhaps augmented by store-operated Ca^2+^ release (Berridge, 1998; Rose and Konnerth, 2001), promotes the movement and fusion of CCK-containing vesicles to the plasma membrane by a syt7-dependent mechanism (Tawfik et al., 2021).

### Somatodendritic release of dopamine and CCK may occur independently

Theoretically, LDCVs that contain DA could also contain CCK. Somatodendritic release of both CCK and DA share some properties, but there also appear to be differences. CCK release was clearly able to potentiate inhibitory synapses after depolarization of a single dopamine neuron voltage-clamped to -40mV (without evoking action potentials) (Figure 1A-C), while somatodendritic release-dependent D2-IPSCs are tetrodotoxin-sensitive (Beckstead *et al*., 2004). The Ca^2+^ chelator, BAPTA, was reported not to block the D2-IPSC, but did prevent depolarization-induced LTP (Figure S3A-C)(Beckstead and Williams, 2007; St Laurent *et al*., 2020). Furthermore, our data suggest that CCK release is blocked by 50 μM Ni^2+^, while somatodendritic DA release was relatively unaffected by T-type channel block (Chen et al., 2006; Ford et al., 2007). Finally, if the trains used to evoke D2 IPSCs had caused the release of CCK, GABA_A_ IPSCs should have undergone LTP, preventing further potentiation upon CCK bath application (Figure 3). The evidence therefore suggests that somatodendritic release of DA and CCK can occur independently.

### Somatodendritic release of dopamine and CCK act over different time scales

Somatodendritic release of DA hyperpolarizes DA cells, and is thought to provide feedback inhibition after a burst of action potentials on a time scale of milliseconds to seconds (Ford et al., 2010; Hikima et al., 2021). When DA and CCK somatodendritic release occur simultaneously, they will act synergistically to inhibit DA neurons, but over different time courses. CCK, by contrast, initiates a gradual, far slower-onset LTP of GABAergic afferents over many minutes, and the synaptic potentiation lasts for over an hour. Synapses on CCK-expressing DA cells may be those most susceptible to modulation by local CCK release, but as neuropeptides can diffuse relatively long distances before being degraded, modulation by CCK may not be a localized event, unlike somatodendritically released dopamine which is expected to undergo rapid reuptake.

### CCK modulates VTA local circuit

Early studies in rats began exploring the effect of CCK on DA neurons and found that it potentiates dopamine-induced inhibition of DA cell firing *in vitro* and *in vivo* (Artaud *et al*., 1989; Brodie and Dunwiddie, 1987). Our data demonstrate that CCK potentiates GABAergic inputs to VTA DA cells after persistent activation of these neurons, providing a brake on their activity, much as CCK released upon repetitive depolarization potentiates GABAergic synapses in the dorsomedial hypothalamus (Crosby *et al*., 2015; Crosby *et al*., 2018). Rather than upregulating all inhibitory synapses, depolarization does not potentiate synapses from a major GABAergic input to the VTA, the RMTg (Jhou et al., 2009; Markovic et al., 2021), but does potentiate those from the ventrolateral periaqueductal gray (St Laurent *et al*., 2020). Similarly, exogenous CCK, or depolarization that releases CCK, selectively potentiates only caudally-evoked but not rostrally-evoked IPSCs (Figure 2E-J)(Dacher and Nugent, 2011)). The simplest way to envision this selectivity is if CCK does not exert autocrine effects on DA cells but instead acts on specific susceptible GABAergic terminals that express the CCK2R. Our FISH data support this idea, as they indicate very low levels of CCK2Rs on somata in the VTA (Figure 2D).

### CCK infused into the VTA decreases food consumption and DA cell activity

What is the role of CCK in VTA DA neurons? Dopamine neurons fire during consumption of food and water and are essential for motivation for food rewards (van Zessen et al., 2012; Zhou and Palmiter, 1995), although they may not be necessary for feeding *per se* (Boekhoudt et al., 2017; Mikhailova et al., 2016; Salamone and Correa, 2012; Sandhu et al., 2018). We found that after an overnight fast bilateral CCK infusion into the mouse VTA blunted food-evoked Ca^2+^ activity with a concomitant reduction in food intake. Thus, CCK may inhibit DA cell activity and reduce feeding in a state-dependent manner, such as during a strongly driven appetitive motivational state (hunger).

As described above, VTA DA neurons release CCK as a negative feedback modulator to tune down their own activity. Upon feeding in the hungry state, DA neurons are engaged to drive the motivation to approach and consume food. However, when caloric intake has reached sufficient levels, DA neurons must be inhibited to reduce the motivation to eat and prevent unneeded intake. Somatodendritic release of CCK is an attractive candidate to mediate this process. Previous work has shown that insulin, which is elevated in the plasma after feeding, depresses excitatory synapses on DA neurons (Labouebe et al., 2013; Liu et al., 2016), and similarly, leptin reduces VTA DA firing rates and suppresses food intake (Hommel et al., 2006). If CCK is released upon feeding, it may act synergistically with insulin and leptin to promote DA cell inhibition and suppress the motivation to continue feeding, consistent with a role in meal termination.

Peptide signaling adds complexity and richness to neural circuits. We propose that the release of neuropeptides from somatodendritic sites may be a common feature of peptide signaling. Our data show that a neuropeptide can be released somatodendritically with relatively modest stimuli but may exert persistent effects on synaptic strength.

## Acknowledgments

The authors would like to thank Dr. Elizabeth Steinberg for helpful discussions, and Dr. Neir Eshel and the STAAR Lab at Stanford for support with fiber photometry experiments. Funding support was from: NIH grants R01 DA011289 (JAK) and T32 DA035165 (MPB), and Stanford Dean’s award (VMD).

## Author contribution

VMD contributed to conceptualization, data curation, formal analysis, investigation, methodology, and writing of the original draft. MBP contributed to conceptualization, data curation, formal analysis, investigation, methodology, and writing. CC and AMW contributed to data curation and formal analysis. RCM contributed to funding acquisition, supervision, and writing of the original draft. JAK contributed to conceptualization, funding acquisition, project administration, supervision and writing.

## Materials availability

This study did not generate new unique reagents.

## STAR METHODS

### Animals

All procedures were carried out in accordance with the guidelines of the National Institutes of Health for animal care and use and were approved by the Administrative Panel on Laboratory Animal Care from Stanford University. This study used Pitx3:GFP (Zhao et al., 2004) (Jackson Laboratory, stock number: 41479-JAX, strain code: B6.129P2-Pitx3tm1Mli/Mmjax), Dat-IRES-Cre (Jackson Laboratory, stock number: 006660, strain code: B6.SJL-Slc6a3tm1.1(cre)Bkmn/J), Ai32 (Jackson Laboratory, stock number: 012569, strain code: B6;129S-Gt(ROSA)26Sortm32 (CAG-COP4*H134R/EYFP) Hze/J), sytVII KO (Jackson Laboratory, stock number: 004950, strain code: B6.129S1-Syt7tm1Nan/J) and C57BL/6 male and female mice bred in-house. Mice were maintained on a 12-h light/dark cycle and provided food and water *ad libitum* except as noted.

### Preparation of brain slices

Mice were deeply anesthetized with ketamine (75 mg/kg) and dexmedetomidine (0.25mg/kg) and quickly perfused with an ice-cold solution containing (in mM): 110 choline chloride; 25 glucose; 25 NaHCO_3_; 7 MgCl_2_; 11.6 sodium ascorbate; 3.1 sodium pyruvate; 2.5 KCl; 1.25 NaH_2_PO_4_ and 0.5 CaCl_2_ saturated with 95%O_2_/5% CO_2_ (pH 7.4; 290-300 mOsm). Horizontal midbrain slices containing the VTA were cut at a thickness of 220 μm using a vibratome (Leica Microsystems), transferred to a holding chamber where the slices were submerged in 37°C warmed artificial cerebrospinal fluid (aCSF) containing (in mM): 126 NaCl, 21.4 NaHCO_3_, 2.5 KCl, 1.2NaH_2_PO_4_, 2.4 CaCl_2_, 1.2 MgSO_4_, and 11.1 glucose, saturated with 95%O_2_/5% CO_2_ (pH 7.4; 290-300 mOsm) for 10 min, and then held at room temperature until transferred to the recording chamber.

### Electrophysiology

Midbrain slices were continuously perfused at 1.5–2 ml/min with aCSF at 28–32°C containing: 6,7-dinitroquinoxaline-2,3-dione (DNQX; Tocris; 10μM), (2R)-amino-5-phosphonopentanoate (APV; Tocris; 100 µM) and strychnine (Sigma-Aldrich; 1μM) to block AMPA-, NMDA- and glycine receptors, respectively. Patch pipettes (2-5 MΩ) were filled with (Sigma-Aldrich; in mM): 125 KCl, 2.8 NaCl, 2 MgCl_2_, 2 ATP-Na+, 0.3 GTP, 0.6 EGTA, and 10 HEPES (pH=7.25-7.28; 265-280 mOsm; junction potential at 30°C 3.4 mV and data are not corrected for the junction potential). In some experiments, 30 mM BAPTA was added to the pipette solution and cells were held for at least 20 minutes prior to depolarization to optimize drug delivery. Whole-cell patch-clamp recordings were made from neurons visually identified in the lateral VTA, recognized in horizontal slices as lateral and rostral to the medial lemniscus and medial to the medial terminal nucleus of the accessory optic tract (MT).

The majority of reported brain slice experiments used Pitx3-GFP mice to label dopamine neurons for identification. In Figures 1A-C and Figures S3G-I, dopamine neurons were identified by the presence of a large I_h_-current (>25 pA) during a voltage step from −50 mV to −100 mV (Ungless and Grace, 2012, Baimel et al., 2017). Expression of I_h_ alone does not unequivocally identify dopamine cells in VTA slices (Margolis et al., 2006). We followed this criterion aware that, in these experiments, a subset of the neurons recorded from and reported here are possibly non-dopaminergic neurons.

DA cells were voltage-clamped at -70 mV except when depolarized to -40mV for 6 min (DEP). Cell input resistance and series resistance were monitored throughout the experiment by means of a -10 mV hyperpolarizing step from -70 mV for 100 ms and cells were discarded if these values changed by more than 15% during the experiment. IPSCs were amplified using an AM Systems amplifier, low-pass filtered at 2 kHz and digitally sampled to computer at 10 kHz using an analog-to-digital interface (Molecular Devices).

### Drugs

Drugs were made up as stock solutions in water, and diluted to appropriate concentrations in aCSF. In experiments using CCK bath application, sulfated cholecystokinin octapeptide, the most abundant form of CCK in the brain (Crawley and Corwin, 1994)(CCK-8S; Tocris; 0.1 μM), was included in aCSF. In these experiments, bovine serum albumin (BSA; Sigma-Aldrich; 0.2 g/L) was added to the aCSF to enhance peptide solubility and minimize nonspecific binding; the protease inhibitor amastatin (Sigma-Aldrich; 50 μM) was also included in the aCSF. The CCK2 receptor antagonist, 2-[2-(5-Bromo-1H-indol-3-yl)ethyl]-3-[3-(1-methylethoxy)phenyl]-4-(3H)-quinazoline (LY225910; Tocris; 1 μM), and sulpiride (Tocris; 150nM) were each dissolved in DMSO and diluted in aCSF for a final concentration of DMSO of 0.1%.

### Stimulation protocols

For synaptic stimulation, a bipolar stainless-steel stimulating electrode was placed caudal to the VTA 200-500 μm from the recorded cell, unless otherwise noted. Rostrally-evoked IPSCs had the stimulating electrode rostral to the cell within the VTA. GABA_A_ receptor-mediated IPSCs were evoked at 0.1 Hz using 100 μsec current pulses. Our stimulation protocol did not produce action potentials escaping voltage clamp, but in the rare cases when cells began firing later in the recording, they were excluded from analysis. For plasticity experiments, after a stable 10 min baseline, the recorded neuron was transiently voltage-clamped to -40 mV for 6 min (DEP) and, only where noted, in some experiments simultaneous synaptic stimulation was delivered at 1 Hz (DEP+LFS). For optogenetic stimulation we delivered light from a white LED (Mightex) controlled by driver (ThorLabs) and reflected through a 40x water immersion lens. For optogenetic stimulation of VTA DA neurons in slices from Ai32::Dat-Cre mice, trains of light (20 Hz, 1000 ms) were delivered for 6 minutes in current clamp mode.

### Analysis of electrophysiological data

IPSC peak amplitudes were measured and responses were normalized by taking the mean of the 10-min baseline responses and dividing the rest of the responses by this mean in GraphPad Prism. These normalized values were then used for average plots. For these plots, all cells were time-aligned at the beginning of the 10-min baseline and averaged over the entire period. The expression of LTP was determined comparing averaged IPSC amplitudes for the 5 min period just before DEP or CCK application with averaged IPSC amplitudes during the 5 min period at 25-30 min after each manipulation.

### Stereotaxic surgeries for viral injections and cannulation

Stereotaxic surgeries were performed on male and female mice between postnatal days 25-35 under deep isofluorane anesthesia (3-5% for induction, 1-2% for maintenance). The mouse’s skull was fixed with ear bars into a stereotaxic apparatus (David Kopf Instruments, Tujunga, CA, USA) with the animal positioned on a warming pad and a craniotomy was drilled at specific coordinates. For viral infusions in VTA for ELISA assays, 200-300 nL of AAV-DJ-EF1a-DIO-hChR2(H134R)-EYFP was injected bilaterally into the VTA of Dat-IRES-Cre mice (mm to Bregma: AP:−2.9, ML: ±0.75, DV: −4.6 from the skull). For fiber photometry recordings, 1 uL of AAV9-FLEX-jGCaMP8m was injected unilaterally into the VTA Dat-IRES-Cre mice (AP: -2.9, ML: -0.75, DV: -4.6 from the skull). The injector pipettes were left in place for 10 minutes before they were slowly retracted. For intracranial cannula placements, a 26-guage bilateral guide cannula (P1 Technologies) was implanted over the VTA (AP: -2.9, ML: ± 0.75, DV -3.2) in wild type mice. Injectors extend 1.5 mm past the guide cannulas, creating a final DV coordinate of -4.7. Cannulas were secured to the skull with stainless steel screws (thread size 00-09 x 1/16, Antrin Miniature Specialties), C&B Metabond, and light-cured dental adhesive cement (Geristore A&B paste, DenMat). Finally, a dummy cannula was inserted into the guide cannula and a dust cap placed to cover and secure the dummy cannula in place. Animals were treated with subcutaneous carprofen 5 mg/kg and topical 2% lidocaine during the surgery for pain control and were allowed to recover for at least 5 days prior to behavioral experiments.

### Immunohistochemistry

Wild type mice were injected with colchicine into the lateral ventricles to limit peptide diffusion (1.0 μl of 7 μg/ml solution). 24 h later mice were anesthetized with ketamine and xylazine (100 and 10 mg/kg, i.p.) and transcardially perfused with 5 ml 1x phosphate buffered saline (PBS), followed by 20 ml paraformaldehyde (PFA; 4%, in PBS). Brains were rapidly extracted, postfixed overnight (12–18h) in 4% PFA at 4°C, and cryoprotected for 48–96h at 4°C in sucrose solution (30% sucrose in PBS) until they sank. Brains were sliced in 50 μm coronal sections, collected consecutively in 24-well plates containing PBS, covered in light-protective material, and stored at 4°C until immunohistochemical processing.

We used the following antibodies: rabbit anti-CCK (1:250, Immunostar 20078, i.c.v. colchicine treatment required), goat anti-GFP, donkey anti-goat Alexa 488 (1:500, Abcam), donkey anti-rabbit Alexa 647 (1:500, Abcam). Sections were washed in PBS 3 times for 10 min (room temperature, shaker), then washed 3 times for 10 min with warmed 5% Triton-X 100 in 1X PBS (room temperature, shaker), and incubated in a blocking solution containing 5% normal donkey serum and 5% BSA in PBS for 1 hr (room temperature, shaker). Next, we incubated slices in primary antibodies overnight for 12–18h at 4° C, in 4% BSA/PBST block solution (PBST: 1X PBS: 9.5 mL, Triton-X: 10µL, BSA: 0.1g, normal donkey serum: 0.5 mL).

After three 10-min PBS washes, sections were again incubated in blocking solution for 1hr, and then incubated for an additional 3 h in blocking solution containing secondary antibodies. After three final 10-min PBS washes, sections were mounted onto glass slides and coverslipped with Fluoroshield containing DAPI Mounting Media (Vectashield). Images were collected on a Zeiss LSM 710 confocal microscope using Nikon software, and minimally processed using FIJI/ImageJ (Schindelin et al., 2012) to enhance brightness and contrast for optimal representation. Fluorescently labeled cells were counted manually.

### ELISA assay

Dat-Cre mice were injected with AAV-DJ-EF1a-DIO-hChR2(H134R)-EYFP into the VTA and allowed to recover for one week. For each assay, horizontal slices containing the VTA from 5 mice were cut and allowed to recover as described above for electrophysiological experiments. At least one hour after slice preparation, the slices were placed in a 24-well plate with 300 µl of aCSF/well. After 5 minutes, the supernatant was collected (control) and replaced by fresh 300 µl of aCSF. 2 s of full-field light pulses were delivered every 15 s for 6 minutes from a blue LED controlled by a master 8 timer (AMPI), and the supernatant (light) collected. CCK concentration in these samples was assessed with a Human/Mouse/Rat CCK Enzyme Immunoassay Kit from RayBiotech (Catalog #: EIA-CCK, EIAM-CCK, EIAR-CCK) and a microplate reader.

### Fluorescence *in situ* hybridization (FISH)

For examination of gene expression in the VTA, coronal sections were processed for fluorescence *in situ* hybridization (FISH) with RNAscope according to manufacturer’s guidelines. Transcripts examined were *Slc6a3 (Dat)* (ACDBio cat# 315441), *Cck* (ACDBio cat# 402271), *Cck1r* (ACDBio cat# 313751), and *Cck2r* (ACDBio cat# 439121). Hybridization was performed using RNAscope Fluorescent Multiplex Kit v2 (Advanced Cell Diagnostics). Slides were cover slipped with Fluoromount-G with DAPI (Southern Biotech, 0100-20) and stored at 4°C in the dark before imaging. Cells were counted and co-localization was measured using a custom macro in Fiji.

### Behavioral tests

#### Food intake

Between 1-6 weeks after surgery for cannula implant, mice were single housed 3 days before starting the experiment and fed ad libitum with regular chow. To maximize motivation to consume food, individually housed mice were fasted overnight starting at the onset of the dark cycle by removing the chow diet from the home cages. The following morning mice receive local intra VTA delivery of either vehicle or CCK (1000pmol/0.5 ul). Immediately afterwards, mice were given access to a pre-weighed food pellet for 1 h. All mice were provided with ad libitum regular chow after the test. At least 5 days after this test, the same mice were tested again with the alternate compound, mice were administered saline or CCK in a counterbalanced, crossover design.

#### Open field

Mice received intra-VTA delivery of CCK or vehicle through the implanted cannula and were immediately placed in square polycarbonate cages (35X35 cm) for a period of 30 min. Mice were administered saline or CCK in a counterbalanced, crossover design. Open-field locomotion was recorded and total distance traveled and time spent in the center were analyzed using video-tracking software (Biobserve Viewer III).

### Fiber photometry

Fiber photometry was performed as previously described (Heifets et al., 2019; Wu et al., 2021) DAT-IRES-Cre mice were unilaterally injected with AAV9-FLEX-jGCaMP8m (Addgene) into the VTA and implanted with optical fiber multiple fluid injection cannulas (AP: -2.9, ML: -0.75, DV: -4.6 from skull) (OmFC, 400 um core, 0.66 NA – Doric Lenses). After 3 weeks recovery, mice were trained to eat palatable fruit loops in the home cage for 1 week. Mice were then habituated to microinjection and photometry procedures. The OmFC implant allows for drug microinjections into the same site being recorded. Injectors (FI_OmFC – Doric Lenses) were secured to a pump-mounted Hamilton syringe with polyethylene tubing (Doric Lenses) and backfilled with sterile H_2_O. Saline or CCK (1000pmol/0.5 ul) was infused into the VTA at a rate of 150nL/min in the home cage. After 1 min of diffusion, injectors were slowly retracted. Mice were then immediately transferred to the fiber photometry room for recordings.

To maximize motivation to consume fruit loops, mice were food deprived overnight prior to each recording day. On recording days, immediately after microinjections, mice were transferred to the recording rooms and connected to patch cables routed through a pigtailed fiber optic rotary joint commutator (FRJ_1×1_PT). After a 5 min baseline, ∼2 grams of fruit loops were placed in the cage and mice were allowed to consume freely for 30 min. Fruit loops were weighed before and after each session. About one week later, mice were subjected through an identical procedure, with the only exception being the injected solution. Mice were administered saline or CCK in a counterbalanced, crossover design.

Data were acquired using Synapse software controlling an RZ5P lock-in amplifier (Tucker-Davis Technologies). GCaMP8m was excited by frequency-modulated 473 and 405 nm LEDs (Doric Lenses). Optical signals were band-pass-filtered with a fluorescence mini cube (Doric) and signals were digitized at 6 kHz. Signal processing was performed with custom scripts in MATLAB (MathWorks). Signals were de bleached by fitting with a mono-exponential, bi-exponential, or cubic polynomial decay function and the ΔF/F signal was z-scored. Videos were manually analyzed and the first 10 consumption events were determined. The analysis was capped at 10 events due to the prediction that CCK suppresses food intake. Peristimulus time histograms were constructed by taking the average of 15 sec epochs of fluorescence consisting of 5 sec before and 10 sec after fruit loop consumption, which is defined as time = 0. Before averaging, each epoch was offset such that the z-score averaged from -5 to -1 sec equaled 0. Peak z-scored fluorescence was determined for each peristimulus time histogram as the maximal z-score value between 0 and +10 sec. A correlation analysis of the differences of z-scores and fruit loop consumption between saline- and CCK-treated sessions was performed. Peak z-score and correlation statistics were calculated with Prism 9.

### Statistics

All data are presented as mean ± standard error. We used GraphPad Prism for statistical analysis. Data were first tested for compliance with parametric testing requirements (normality and homoscedasticity). Parametric and nonparametric data were then analyzed using a paired t-test or a Wilcoxon test for significance. Differences were deemed to be significant with p values <0.05.

---

**Supplementary Figure 1.**
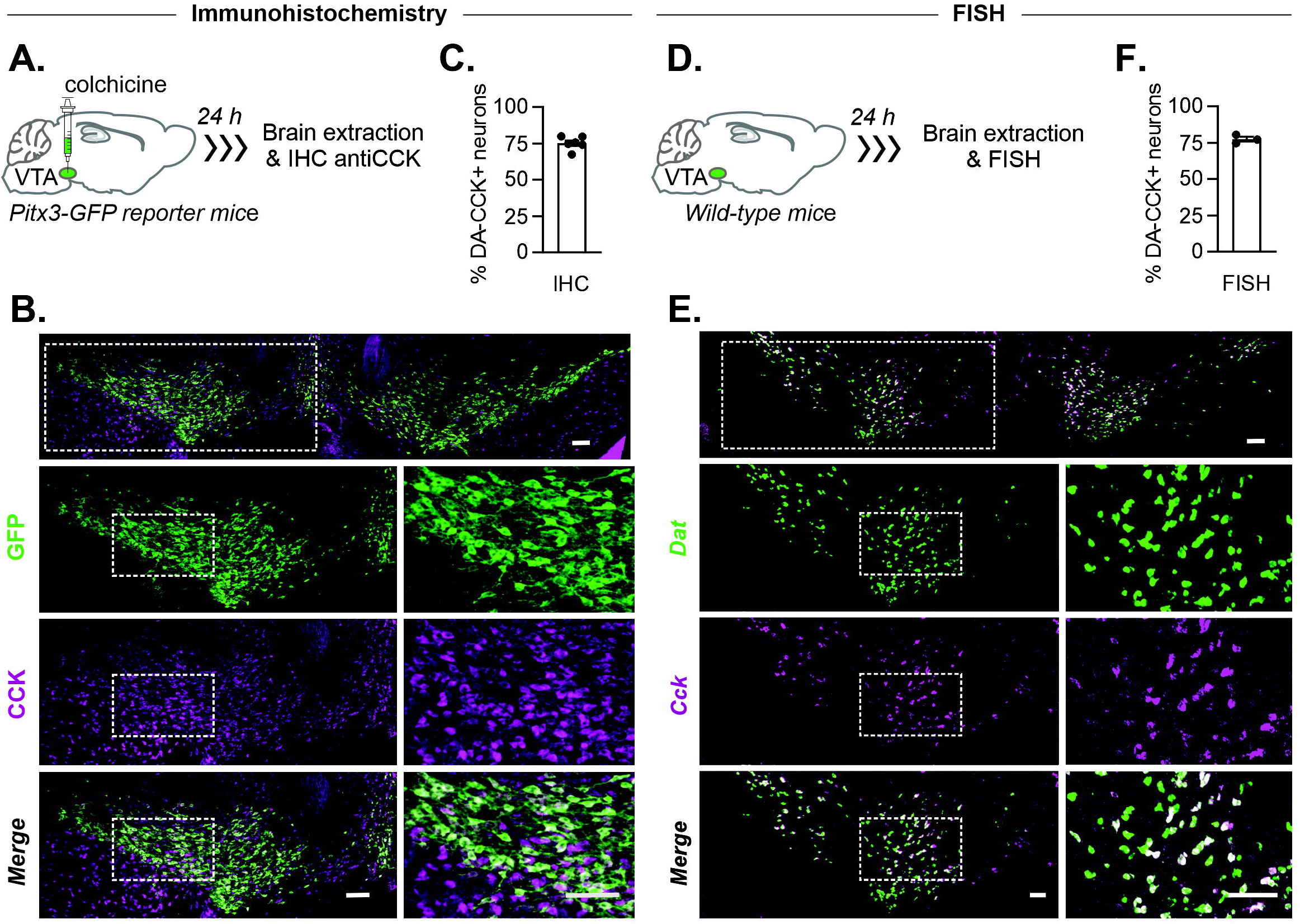
CCK is expressed in the majority of DA neurons in the VTA. **A.** Diagram of the strategy used for immunohistochemistry, colchicine injection in the VTA of PITX3-GFP reporter mice. **B.** Representative confocal images of fluorescence in a midbrain coronal section containing the VTA. ‘GFP’: GFP signal from Pitx3-GFP (DA) neurons, ‘CCK’: CCK immunostaining, and ‘Merge’: overlay of GFP and CCK. Scale bar=10 µm. **C.** Bar graph indicating percentage of GFP+ neurons that co-express CCK using immunohistochemistry. **D.** Representative confocal images of a FISH experiment on a coronal slice containing the VTA. ‘DAT: signal from DA transporter (*Slc6a3*)-expressing neurons, CCK, and ‘Merge’: overlay of DAT and CCK. Scale bar=10 µm **E.** Bar graph indicating percentage of DAT+ neurons that co-express CCK using FISH.

**Supplementary Figure 2.**
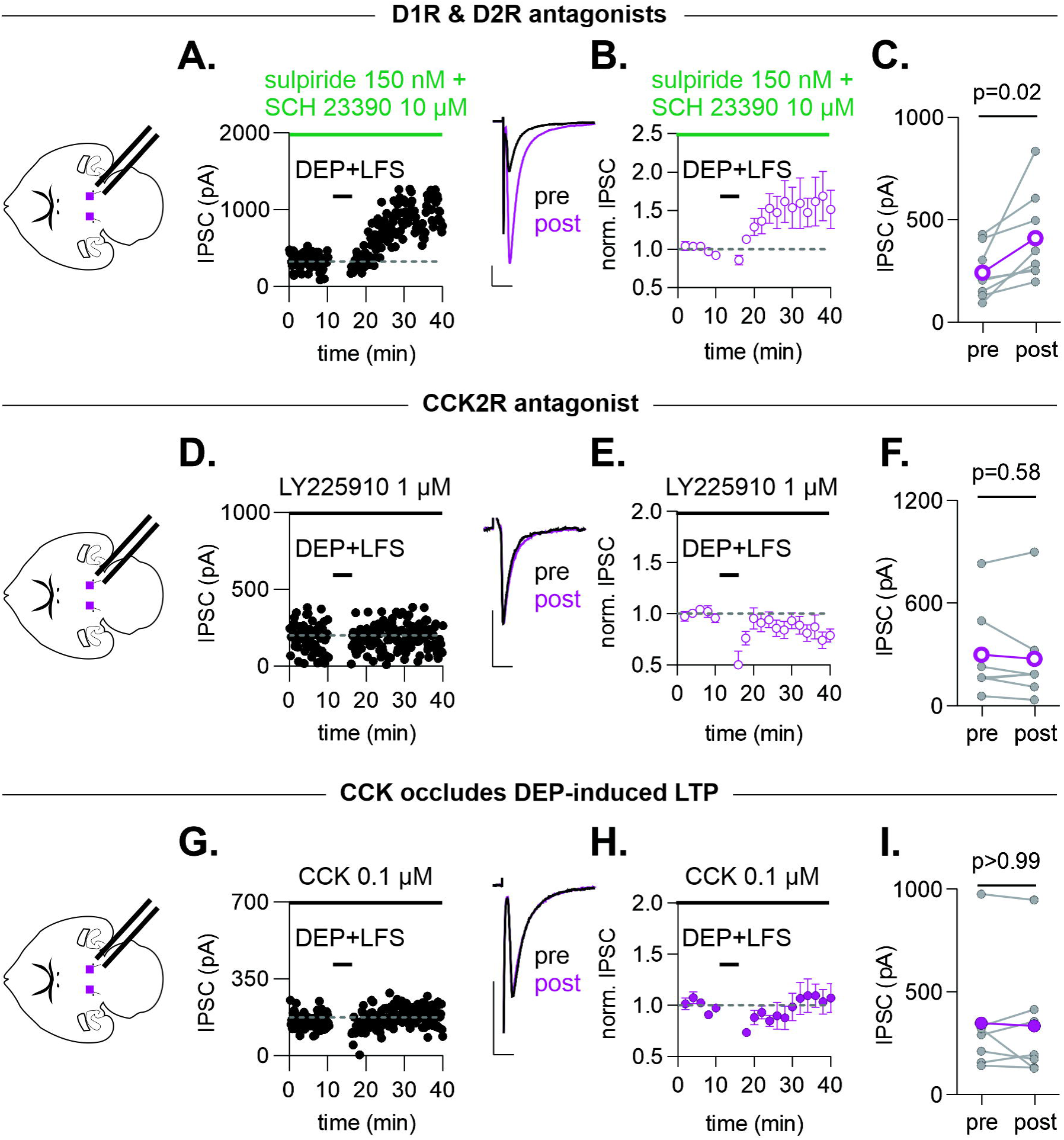
CCK and not dopamine mediates LTP evoked by depolarization plus LFS. **A.** Representative time course and example IPSCs before and after DEP+LFS in the presence of sulpiride (150 nM) and SCH 23390 (10 µM). **B.** Time course of averaged IPSC amplitudes before and after DEP+LFS (n=8 cells/4 mice). **C.** IPSC amplitudes before (5-10 min) and (25-30 min) after DEP+LFS (n=8 cells/4 mice). Paired t-test, p=0.02. **D.** Representative time course and example IPSCs before and after a 6-minute depolarization of the recorded neuron from -70 to -40 mV with simultaneous afferent low frequency (1 Hz) stimulation (DEP+LFS) in the presence of LY225910 (1 µM). **E.** Time course of averaged IPSC amplitudes before and after DEP+LFS (n=7 cells/4 mice). Colored symbols/lines represent the mean response across all cells, error bars represent SEM. **F.** IPSC amplitudes before and after DEP+LFS (n=7cells /4 mice). Wilcoxon t-test, p=0.58. **G.** Representative time course and example IPSCs before and after (DEP+LFS) in the presence of CCK. **H.** Time course of averaged IPSC amplitudes before and after DEP+LFS (n=7 cells/5 mice). **I.** IPSC amplitudes before and after DEP+LFS (n=7 cells/5 mice). Wilcoxon t-test, p>0.999. Colored symbols/lines represent the mean response across all cells, error bars represent SEM. Scale bars, 100 pA and 10 ms.

**Supplementary Figure 3.**
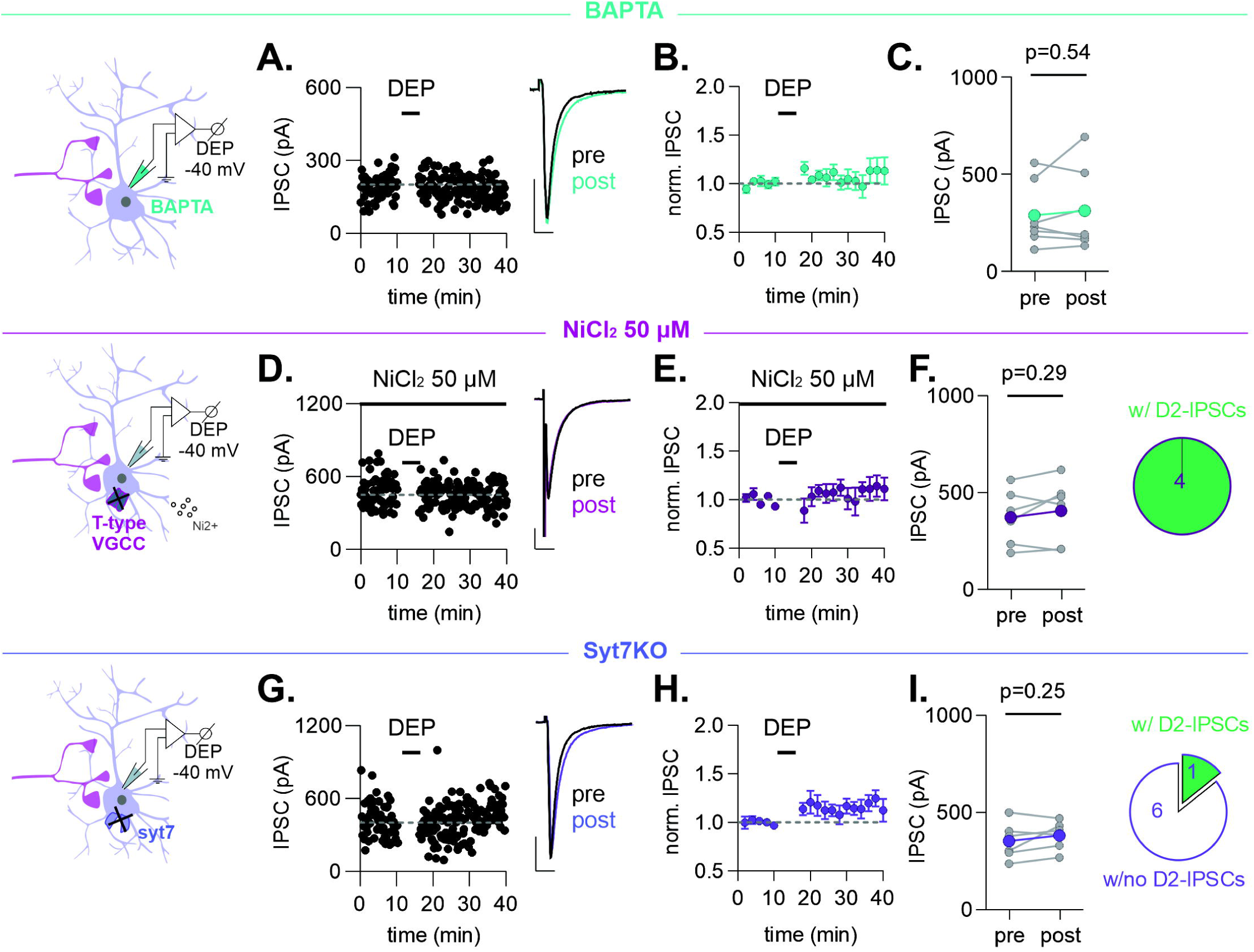
Mechanisms underlying somatodendritic depolarization-induced CCK release. **A.** Representative time course and example IPSCs before and after a 6-minute depolarization of the recorded neuron from -70 to -40 mV (DEP) with BAPTA (30 mM) included in the patch pipette. Recordings were initiated after 20 or more minutes in whole-cell configuration to allow the BAPTA to diffuse into the cell. **B.** Time course of averaged IPSC amplitudes before and after DEP (n=7 cells/4 mice). **C.** IPSC amplitudes before and after DEP (n=7 cells/4 mice). Paired t-test, P=0.54. **D.** Representative time course and example IPSCs before (pre) and after (post) DEP in the presence of NiCl_2_ (50 µM). **E.** Time course of averaged IPSC amplitudes before and after DEP (n=7 cells/3 mice). **F.** IPSC amplitudes before and after DEP (n=7 cells/3 mice). Paired t-test, p=0.29. Pie chart indicates that in 4/4 neurons, D2-IPSCs could be evoked. **G.** Representative time course and example IPSCs before and after DEP in a VTA dopamine neuron from a syt7KO mouse. **H.** Time course of averaged IPSC amplitudes before and after DEP in dopamine neurons from syt7 KO mice (n=6 cells/3 mice). Pie chart indicates that in 6/7 syt7-KO neurons, D2-IPSCs could not be evoked. **I.** IPSC amplitudes before and after DEP (n=6 cells/3 mice). Paired t-test, p=0.25. Scale bars represent 100 pA and 10 ms.

**Supplementary Figure 4.**
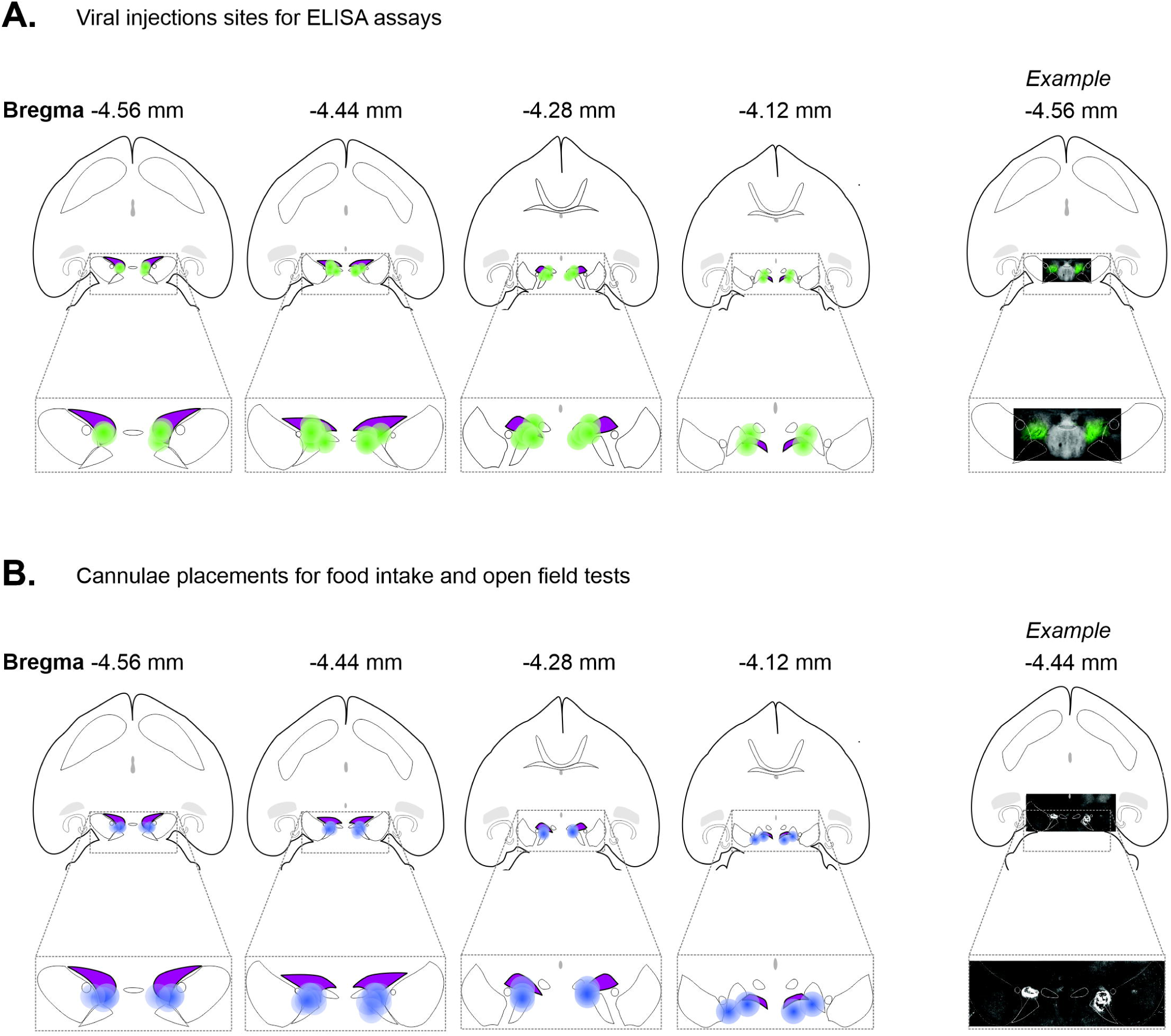
Viral injection sites and cannulae placements in infusion experiments. **A.** Schematic of horizontal brain slices indicating viral injection sites in DatCre mice used for ELISA assays. Purple region indicates VTA location and green dots indicate viral injection central area. **B.** Schematic of horizontal brain slices indicating cannulae placements for food intake and open field tests. Purple region indicates VTA location and blue dots indicate bilateral cannulae location.

